# Maternally-inherited anti-sense piRNAs antagonize transposon expression in teleost embryos

**DOI:** 10.1101/2021.11.03.467172

**Authors:** Yixuan Guo, Krista R. Gert, Svetlana Lebedeva, Magdalena E. Potok, Candice L. Wike, Edward J. Grow, René F. Ketting, Andrea Pauli, Bradley R. Cairns

## Abstract

Transposable elements threaten genome stability, and the Piwi-piRNA system has evolved to silence transposons in the germline^1–6^. However, it remains largely unknown what mechanisms are utilized in early vertebrate embryos prior to germline establishment and ‘ping-pong’ piRNA production. To address this, we first characterized small RNAs in early zebrafish embryos and detected abundant maternally-deposited, Ziwi-associated, antisense piRNAs that map largely to evolutionarily young long terminal repeat (LTR) retrotransposons. Notably, the focal establishment of the repressive modification H3K9me2/3 coincides with these young LTR elements, is deposited independent of transcription, and is required for LTR silencing. We find piRNAs highly enriched and maintained in primordial germ cells (PGCs), which display lower LTR expression than somatic cells. To examine the consequences of piRNA loss, we used reciprocal zebrafish-medaka hybrids, which display selective activation of LTRs that lack maternally-contributed targeting piRNAs. Thus, the Piwi-piRNA system actively antagonizes transposons in the soma and PGCs during early vertebrate embryogenesis.

## Results

To characterize the small non-coding RNA (sRNA) repertoire during zebrafish early embryogenesis and to assess the presence (and inheritance) of piRNAs, we performed sRNA-seq of pre-zygotic genome activation (ZGA) (2 hpf and 2.5 hpf) and post-ZGA (4 hpf, 5.3 hpf, 8 hpf and 24 hpf) embryos, as well as sperm and oocytes, with three biological replicates for each stage. sRNA size distribution revealed abundant piRNA-sized sRNAs (24-32 nt) present in early embryos and oocytes, but not in sperm (Figure 1A and S1B), which is consistent with the absence of Piwi proteins in mature sperm^1^. These maternally-deposited, piRNA-sized sRNAs dominate the pre-ZGA embryo sRNA repertoire (>86%, Figure 1A, right table), which resembles sRNA profile from 1-cell stage ^7^. Following ZGA, an sRNA transition occurs, involving the known activation of the miR-430 locus^8,9^ (Figure 1A).

**Figure 1.**
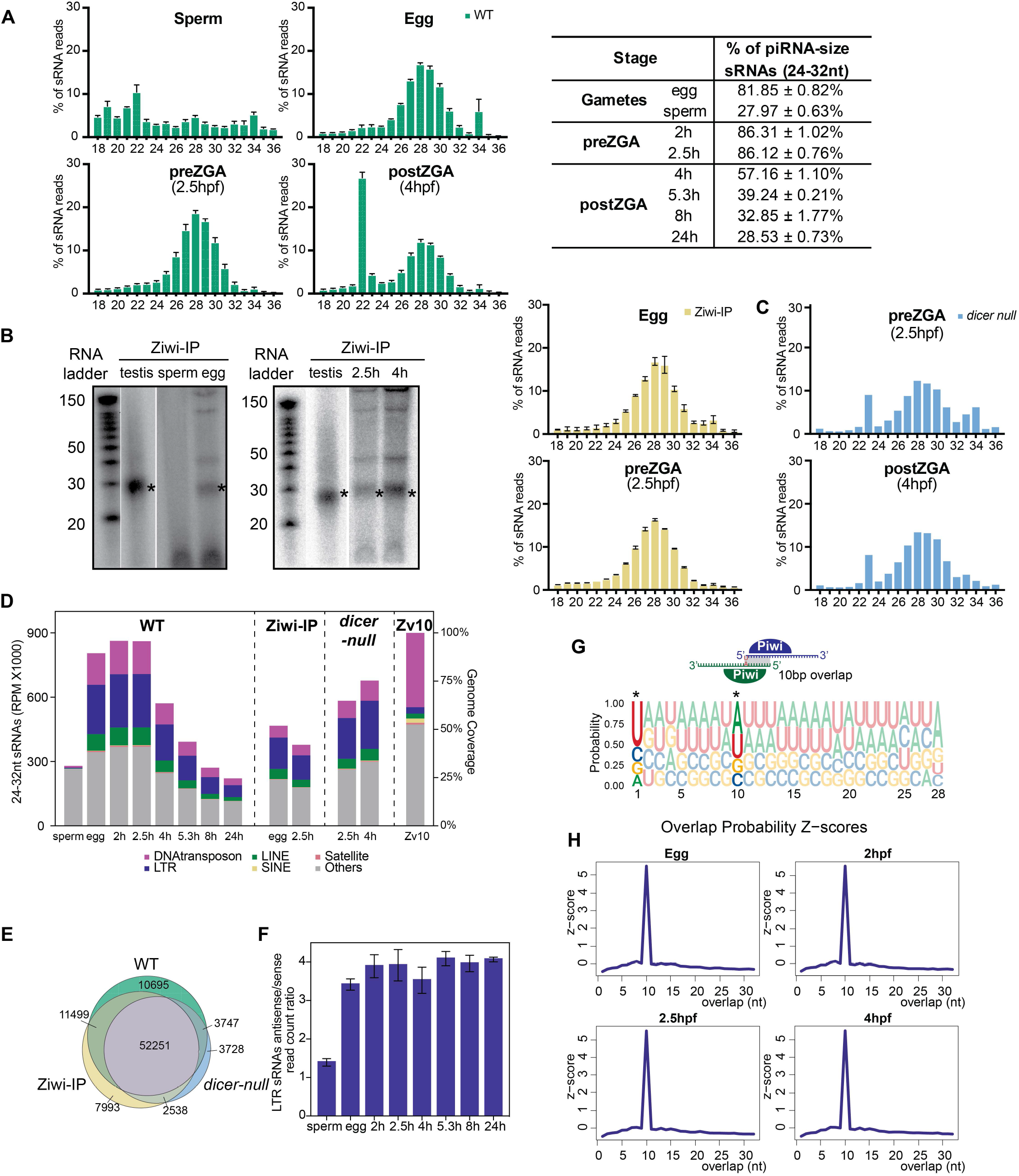
Characterization of the sRNA repertoire in zebrafish gametes and early embryos. **(A)** Left: size distribution of sRNAs in wild-type zebrafish sperm, egg, pre-ZGA (2.5 hpf) and post-ZGA (4 hpf) embryos. Right: quantification of piRNA-size (24-32 nt) sRNAs in zebrafish gametes and early embryos. Data are represented as means± SEM. **(B)** Left: ~30 nt piRNAs are detected from egg and early embryos, but not from sperm. Right: size distribution of piRNAs pulled down by Ziwi antibody from egg and pre-ZGA embryos. **(C)** Size distribution of sRNAs in *dicer-null* embryos. **(D)** Genome distribution of24-32 nt sRNA in WT, Ziwi-IP and *dicer-null* samples. The genome composition of the GRCzlO reference genome is shown on the right. **(E)** The intersection ofLTRs targeted by piRNAs from WT, Ziwi-IP and *dicer-null* samples. LTRs with >= l mapped sRNA reads are included. **(F)** The ratio between antisense and sense LTR-targeting piRNAs from WT gametes and early embryos. **(G)** Top: Schematic representation ofpiRNAs sequence bias and ping-pong signature. Bottom: sRNAs from pre-ZGA embryos (2.5 hpf) carry the sequence bias ofUl and AlO (highlighted and marked by*). **(H)** The ping-pong signature (peak at 10 bp overlap) is detected from the sRNA population in eggs and early embryos.

To test whether these piRNA-sized sRNAs are truly piRNAs, we assessed their physical association with the sole zebrafish Piwi family protein present in the early embryos Ziwi by immunoprecipitation (IP) from zebrafish gametes and early embryos, as Zili protein is absent prior to 3 dpf^7,10^.We found that piRNA-sized sRNA (~30 nt) co-precipitated with Ziwi from oocytes and early embryos, but not sperm (Figure 1B). Furthermore, and consistent with these sRNAs being indeed piRNAs, the piRNA-sized population was not affected in *dicer-null* embryos whereas activation of miRNA expression was largely suppressed (Figure 1C).

Genome mapping of piRNA-sized reads from wild-type oocytes and embryos, Ziwi-IP, and *dicer-null* embryos displayed strong enrichment at repetitive elements, especially at long terminal repeat (LTR) retrotransposons (Figure 1D and 1E). Our oocyte results are consistent with prior sequencing of and Northern analysis of the zebrafish ovary ^1,10^. In contrast, sperm-derived reads lacked LTR enrichment. Notably, reads mapping to LTRs showed a strong (3-4-fold) antisense strand bias (Figure 1F). They also carried a major sequence bias, namely a uracil at the first position (U1), and a minor bias of an adenine at the tenth position (A10) (Figure 1G and S1C). Although there is only one Piwi protein in early zebrafish embryos (Ziwi), we clearly detected the ping-pong signature, characterized by a distinct peak of a 10-bp overlap from embryonic piRNAs (Figure 1H and S1D). As testes and ovaries contain both Zili and Ziwi, and as pre-ZGA embryos do not conduct transcription to generate new substrates, the repertoire of piRNAs in early embryos is evidently generated in oocytes and deposited into early embryos.

In principle, the Piwi-piRNA pathway could mediate transposon silencing post-transcriptionally by targeting complementary RNAs for degradation. Alternatively, it could also function at the transcriptional level by affecting DNA methylation and/or the repressive histone modifications H3K9me2/3. However, DNA and LTR transposons in early zebrafish embryos are DNA hypermethylated^11,12^, which is the ‘default’ state in the zebrafish genome at locations without placeholder nucleosomes (which bear H2A.Z and H3K4me1^13^). Placeholder nucleosomes deter DNA methylation during early embryogenesis by preventing maintenance DNA methyltransferases after replication ^13^, as zebrafish embryos do not conduct genome-wide DNA demethylation after fertilization ^12,14^. Thus, as DNA methylation is constitutive at transposons, an additional heterochromatin mark(s) may be needed for focal repression.

To assess whether piRNAs could in principle impact H3K9 methylation to silence transposons, we first tested whether histone methylation levels were changing during early zebrafish embryogenesis. Immunostaining revealed H3K9me2 and H3K9me3 (Figure 2A and S2A) detectable in 1-cell stage embryos (15 min. post-fertilization) but absent at the 64-cell stage. However, by the 256-cell stage (pre-ZGA), H3K9me2 was largely restored, whereas H3K9me3 was observable only from major ZGA onwards (sphere stage). Transcription inhibition by α-amanitin (treated at the 1-cell stage, leading to late sphere-stage arrest (4 hpf)) conferred only a slight decrease in H3K9me3 and did not affect H3K9me2 (Figure S2A), suggesting that transcription is not necessary for the acquisition of H3K9me2 nor for the *de novo* establishment of H3K9me3.

**Figure 2.**
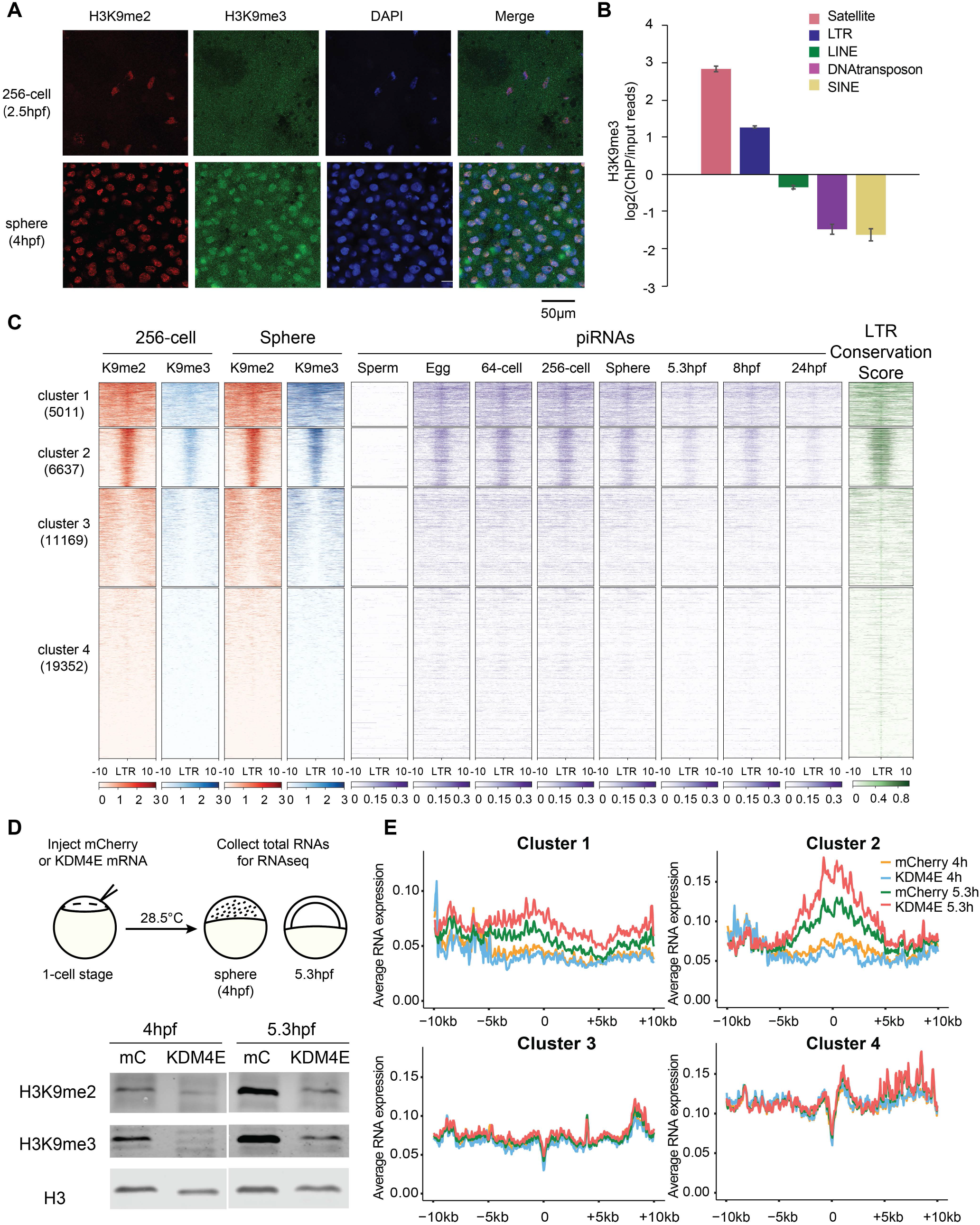
Establishment of the repressive histone marks H3K9me2/3 at piRNA-enriched, evolutionary young LTRs. **(A)** Immunofluo-rescence staining ofH3K9me2 (red), H3K9me3 (green) and DAPI (blue) in pre-ZGA (256-cell stage, 2.5 hpf) and post-ZGA (sphere, 4 hpf) embryos. **(B)** Sphere stage H3K9me3 ChIP-seq reads enrich at satellite repeats and LTR retrotransposons in the genome. **(C)** Heatmap of H3K9me2 (red), H3K9me3 (blue), embryonic piRNAs, and conservation score at LTR internal regions. K-mean clustering shows that LTR internal regions from clusters 1 and 2 have high H3K9me2 pre- and post-ZGA and gain H3K9me3 during ZGA, whereas cluster 3 and 4 lack H3K9me2/3 enrichment. Egg and embryonic piRNAs enriched at clusters 1 and 2. LTR conservation score (green) indicate that clusters 1 and 2 are young elements. **(D)** Top: schematic representation of the experimental procedure ofH3K9me2/3 removal in early embryos. Bottom: both H3K9me2 and H3K9me3 levels are significantly attenuated by KDM4E expression at 4 hpf and 5.3 hpf. **(E)** Metaplots of the LTR internal region expression by cluster in mCherry or KDM4E mRNA injected embryos at 4 hpf and 5.3 hpf. Clusters are defined in panel C.

To localize H3K9me2 and H3K9me3 and to compare to piRNA-mapping regions, we conducted chromatin immunoprecipitation followed by sequencing (ChIP-seq) at pre-ZGA (256-cell stage) and post-ZGA (sphere). Notably, H3K9me2 (at pre-ZGA and post-ZGA) and post-ZGA H3K9me3 coincided in early embryos, whereas very limited overlap was observed in adult tissues ^15^. Genomic mapping revealed H3K9me2/3 enrichment exclusively at two types of repeats, satellite repeats and LTRs (Figure 2B and S2B), consistent with prior work on H3K9me3 ^16^. Since embryonic piRNAs were enriched at LTRs but not at satellite repeats we focused on LTR silencing. We observed robust H3K9me3 deposition at LTR internal regions post-ZGA, but only at regions that were initially marked by H3K9me2 pre-ZGA, specifically at clusters 1 and 2, in a transcription-independent manner (Figure 2C and S2C and D). Remarkably, H3K9me3 and piRNAs from eggs and early embryos robustly and specifically co-enrich within these two clusters. Furthermore, the evolutionary conservation score (calculated based on the divergence, deletions, and insertions of LTRs^17^) supports clusters 1 and 2 as more conserved and thus likely more recent LTR insertions. By contrast, H3K9me2/3-depleted LTRs (clusters 3 and 4), which were less conserved (older) than LTRs in clusters 1 and 2, lacked targeting piRNAs. Analysis of open chromatin via ATAC-seq data revealed that the establishment of H3K9me3 at ZGA was accompanied by decreased chromatin accessibility at clusters 1 and 2 (Figure S2C), which is consistent with their transcriptional silencing.

To test whether loss of H3K9me3 indeed derepresses these LTRs, we expressed KDM4E, a placental animal-specific H3K9 demethylase, in early zebrafish embryos. KDM4E expression greatly reduced H3K9me2/3 levels at 4 hpf and 5.3 hpf (Figure 2D). Interestingly, RNA-seq analysis revealed the selective activation of young LTRs (clusters 1 and 2) at 5.3 hpf, which was further enhanced upon H3K9me3 removal by KDM4E expression. In contrast, older/less conserved LTRs (clusters 3 and 4), which lack H3K9me2/3, were not affected (Figure 2E). Taken together, our results show that evolutionary young LTR elements have high H3K9me3 which is correlated with high levels of antisense piRNAs during ZGA, and that young LTRs are kept silent during ZGA by means of their H3K9 methylation.

Zebrafish primordial germ cells (PGCs) are specified during early embryogenesis through germ plasm inheritance and are restricted to four cells at ZGA^18^. Re-analysis of published cell-type specific RNA-seq datasets from 7 hpf embryos^19^ revealed young LTRs (e.g. cluster 2) specifically expressed in somatic cells but not PGCs (Figure 3A; clusters are defined in Figure 2C). This raised the possibility that LTR chromatin and/or piRNA abundance (for transcriptional and/or post-transcriptional regulation) might differ in PGCs and somatic cells, as may be expected, at least for piRNAs, from the enrichment of Ziwi on the PGC-specifying germ plasm^1^.

**Figure 3.**
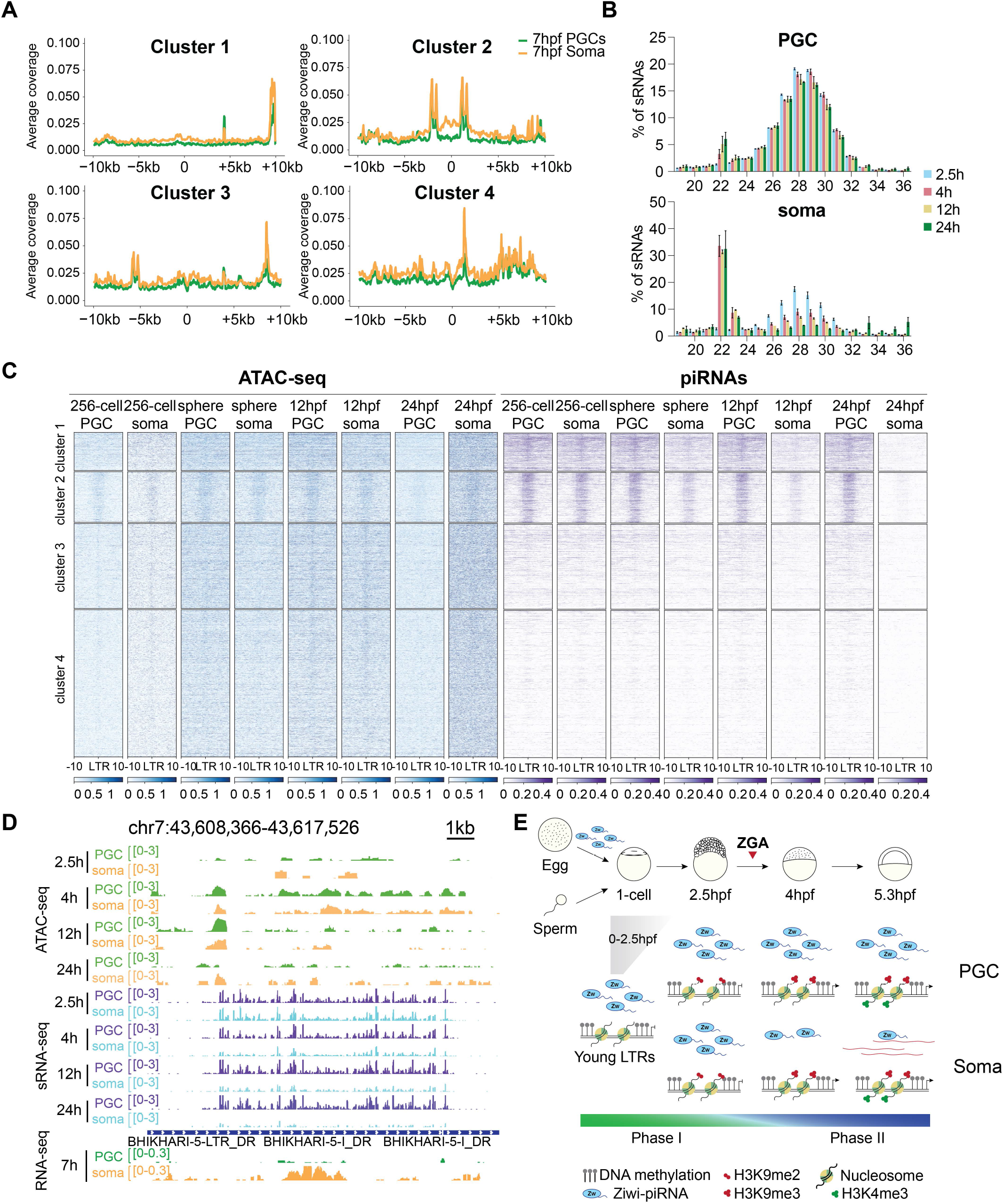
The abundance of piRNAs and transposon transcripts diverges in PGCs and somatic cells. **(A)** Metaplots ofLTR internal region expression by cluster in PGCs (green) and soma (orange) from 7 hpf embryos. **(B)** Size distribution of sRNAs in PGCs and soma from pre-ZGA (2.5 hpf) and post-ZGA (4 hpf, 12 hpf and 24 hpf) embryos reveals maintenance of piRNAs specifically in PGCs. **(C)** The chromatin accessibility and piRNA abundance at LTR internal region clusters in PGCs and somatic cells. Clusters in panel A and C are the same as Figure 2C. **(D)** A genome browser snapshot of representative LTR internal region shows higher abundance of LTR transcripts and lower abundance of piRNAs in soma than in PGCs, but no difference in the extent of chromatin openness/accessibility (by ATAC-seq). **(E)** A model depicting piRNA protection of the genome via transposon silencing in two phases during early embryogenesis. During phase I, maternal piRNAs are deposited into all cells in the early embryo, prior to the establishment ofrepressive marks and prior to germ cell specification. At 2.5 hpf, H3K9me2 is established at piRNA-targeted young LTRs, followed by H3K9me3 during ZGA. The diverging piRNA populations in PGC and soma indicates the beginning of phase II. We hypothesize that piRNAs are specifically retained/maintained in PGCs to continue to keep young LTRs attenuated in germ cells but not in somatic cells.

To examine this, we performed ATAC-seq and sRNA-seq in PGCs and somatic cells at 2.5 hpf, 4 hpf, 12 hpf and 24 hpf. We found similar levels of chromatin openness at LTRs in these two cell types at all time points except for PGCs at 24 hpf, which showed closed chromatin and retained piRNAs at cluster 2 (Figure 3C and S3D). Interestingly, the piRNAs retained in PGCs represent the dominant sRNA type and correspond specifically to conserved young LTRs (clusters 1 and 2; in Figure 3B and C), whereas miRNA-sized (~22 nt) RNAs form only a small peak even after ZGA. In striking contrast, the piRNAs in somatic cells are abundant only prior to ZGA, after which miRNAs dominate and piRNAs gradually diminish. Taken together, these results suggest that piRNAs are maintained selectively in the germline cells

Loss of Piwi-piRNA pathway components Ziwi or Zili causes sterility in zebrafish and prevents sperm or egg production, precluding genetic knockout approaches for examining their functions in early embryos. We therefore utilized zebrafish-medaka hybrids as a valuable tool. Zebrafish and medaka are evolutionarily distant teleost fish that diverged ~110-150 MYA^20^, with largely different TE compositions and thus likely largely different piRNA populations. Both zebrafish-medaka hybrids and the reverse cross can be generated using *in vitro* fertilization^21,22^. These hybrids progress past gastrulation to the segmentation stage or stall at gastrulation (for embryos derived from the reciprocal cross), but do not develop further. Based on our and published data, zebrafish embryos inherit piRNAs exclusively from maternal deposition. Thus, the medaka/zebrafish hybrid system enables functional evaluation in the absence of paternal-genome-matching piRNAs on a hybrid genome with divergent paternal TE targets in both parental orientations.

To analyze piRNA inheritance and LTR activation in the hybrids and purebred fish, we performed sRNA-seq and total RNA-seq for four types of embryos: zebrafish in-cross and zebrafish female to medaka male (termed ‘zebrafish hybrids’, named by the maternal side) at 3 hpf, 5 hpf, 8 hpf and 24 hpf, as well as medaka in-cross and medaka female to zebrafish male embryos (termed ‘medaka hybrids’) at 3 hpf, 5 hpf, and 8 hpf (Figure 4A). The dynamics of sRNA length profile from hybrid embryos resemble the profile of their maternal purebred (Figure 4B). In keeping with their largely unique TE composition, small RNAs from zebrafish and medaka in-cross embryos display minimal overlap. Moreover, as was seen in purebred zebrafish embryos, virtually all LTR-derived piRNAs (24-32 nt) in hybrid embryos were derived from maternal/oocyte inheritance and not from sperm (Figure 4C). Also consistent with the results obtained from zebrafish, medaka and hybrid piRNAs display a sequence bias of U1 and A10, and the ping-pong signature of a peak of a 10-bp overlap (Figure 4E and F). Quite surprisingly, the strand bias of piRNAs was only present in zebrafish and zebrafish hybrids, but not in medaka or medaka hybrids (Figure 4D).

**Figure 4.**
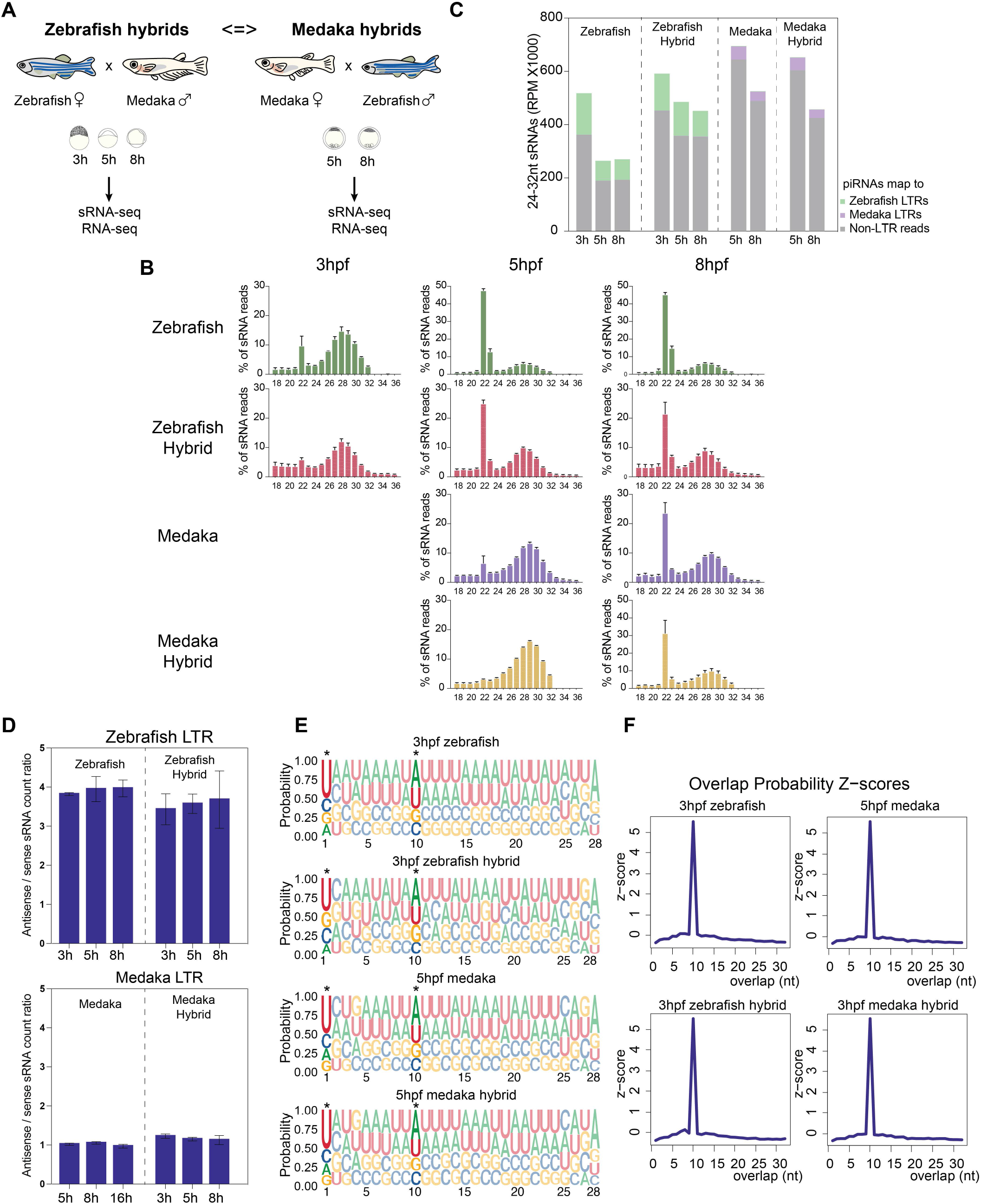
Characterization of sRNA profiles in zebrafish-medaka hybrid embryos. **(A)** Schematic representation of the experimental setup for generating zebrafish and medaka hybrid embryos by reciprocal crosses. Note: hybrid embryos are named by the maternal side. **(B)** Size distribution of sRNAs in zebrafish, medaka, zebrafish hybrid and medaka hybrid embryos. **(C)** LTR-targeting piRNAs in hybrid embryos are primarily contributed by maternal deposition. **(D)** The strand bias ofLTR-targeting sRNAs is only observed in zebrafish and zebrafish hybrid embryos, but not in medaka or medaka hybrid embryos. **(E)** piRNA-size sRNAs in zebrafish, medaka, zebrafish hybrids and medak hybrids carry piRNAs sequence bias ofUl and AlO (highlighted and marked by*). **(F)** piRNA-size sRNAs in zebrafish, medaka, zebrafish hybrids and medaka hybrids carry the piRNA ping-pong signature (1 Obp overlap).

In hybrid embryos, genes from both parental origins are induced during development and hybrid ZGA timing generally aligns with their maternal and not paternal ZGA timing ^22^. To determine whether the absence of paternally-encoded piRNAs in hybrids specifically confers paternal genome LTR activation, we examined the expression of all LTR retrotransposons in our in-cross and hybrid embryos (Figure 5A, S4B and S4C). Indeed, the heatmap of medaka-origin LTRs revealed hundreds of LTRs activated in zebrafish hybrids that are largely attenuated in medaka in-cross embryos. Notably, medaka genome-encoded piRNAs targeting these attenuated LTRs were enriched in medaka in-cross and medaka hybrid embryos, but were not present in either zebrafish in-cross or zebrafish hybrid embryos (Figure 5A, B and S4D), which is consistent with their maternal inheritance. In contrast, medaka hybrids activated hundreds of LTRs of zebrafish/paternal origin, which lacked maternally-deposited targeting piRNAs. LTR expression in the medaka hybrid is less pronounced than in zebrafish hybrids, likely due to the latest (8 hpf) time-point analyzed for medaka hybrid embryos being close to the time-point of ZGA in medaka (note: medaka hybrids are inviable by 16 hpf). Taken together, these results provided by our reciprocal hybrid data causally link maternally-deposited piRNAs to transposon silencing during early embryogenesis (Figure 5C).

**Figure 5.**
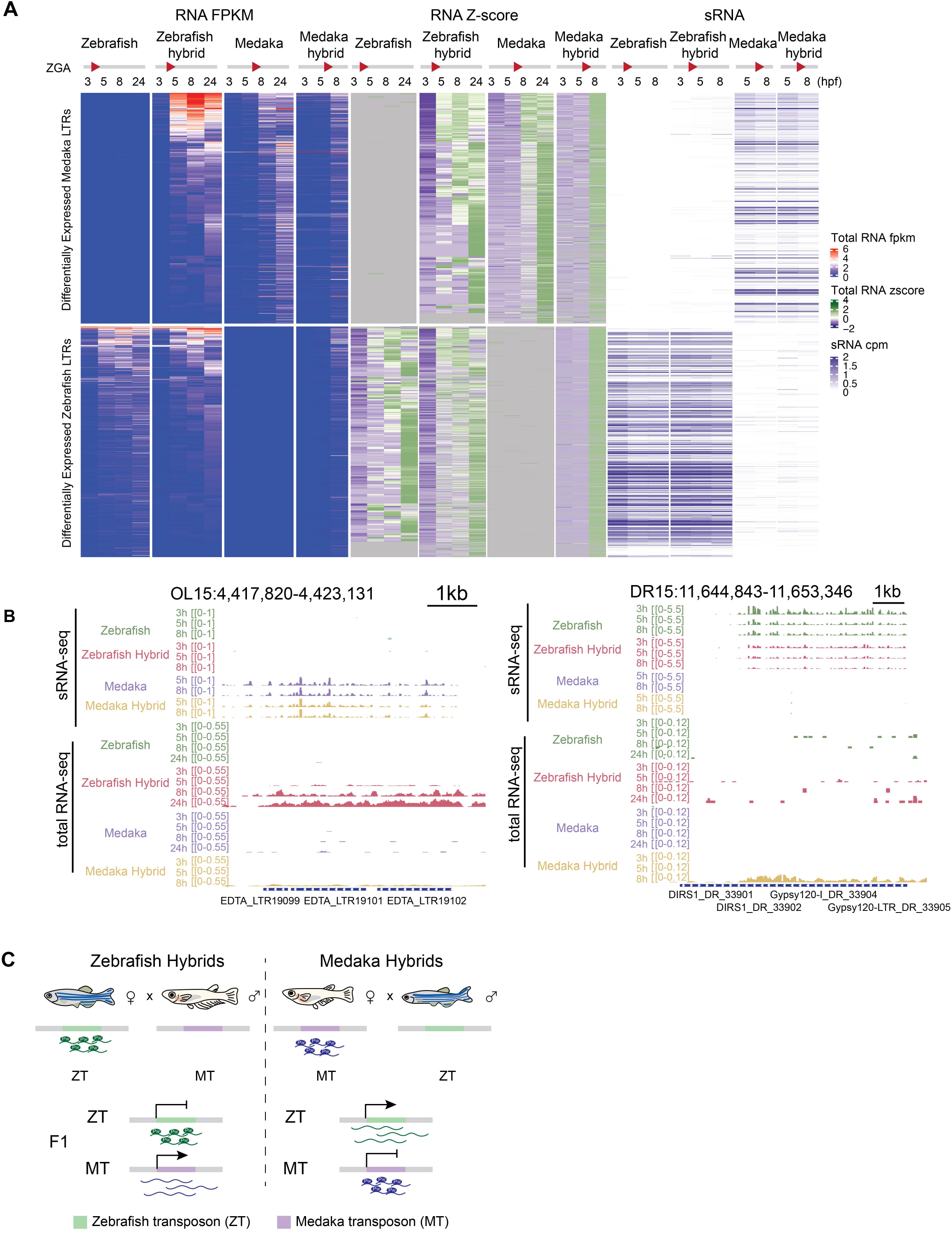
LTRs from the paternal genome are selectively desilenced in the absence of parental-encoded piRNAs. **(A)** Heatmaps of the expression of activated medaka-(top) and zebrafish-(bottom) origin LTRs (FPKM and Z-score) and sRNA abundance in zebrafish, medaka, zebrafish hybrid, and medaka hybrid embryos. ZGA time is marked by the red triangle next to the developmental time. **(B)** Examples of activated medaka (left) and zebrafish (right) LTRs in zebrafish hybrid and medaka hybrid embryos, respectively. **(C)** A graphic summary of the sRNA and total RNA-seq results from hybrid embryos. The absence of piRNAs targeting the paternal LTRs results in their desilencing.

## Discussion

The Piwi-piRNA system plays a central role in transposon silencing in many organisms and operates in the vertebrate germline to protect the integrity of the inherited genome. However, a major gap in our understanding involves the strategy for transposon silencing during early vertebrate embryogenesis—a vulnerable time during which the production of piRNAs (and all transcription) has ceased, heterochromatin is low or absent, and the germline has not yet been defined. Of note, zebrafish lack KRAB-ZNF proteins, which are utilized extensively in mammals for transcriptional silencing of transposons. Here, through a variety of approaches, we explored the notion of parental Piwi-piRNA inheritance and implementation as a solution for transposon silencing in teleost embryos and in germline cells.

Our work provides a series of new findings on piRNA inheritance and function in early embryos and the developing germline. First, early zebrafish and medaka embryos both contain abundant piRNAs, which are exclusively maternally-inherited. In zebrafish, they are primarily anti-sense piRNAs and are solely associated with Ziwi, as Zili protein is absent in early embroys, and only returns during early larval period (3 dpf) ^7^. As Zili is absent, piRNA generation does not occur during early embryogenesis, thus requiring reliance on the maternally-inherited Ziwi-bound piRNAs.

We find that maternally-provided piRNAs are predominantly enriched at evolutionarily young LTR retrotransposons in early embryos, and that during development they are more specifically retained within the PGCs. Later during germ cell development, this maternal piRNA pool has been implicated in fueling the zygotic ping-pong cycle ^7^. Although inherited piRNAs are more efficiently sequestered and maintained in the nascent PGCs than in somatic cells of the embryo, maternal piRNAs are also evident in somatic cells in early embryos. Importantly, we utilized reciprocal zebrafish-medaka hybrids to provide functional evidence consistent with the use of maternally-inherited piRNAs to antagonize LTR transcript levels. Notably, the embryonic piRNAs in zebrafish coincide with locations of initial H3K9me2/3 heterochromatin establishment during early embryogenesis. This deposition of H3K9me2/3 contributes to transcriptional silencing, as diminishing H3K9me2/3 leads to increased transcription from LTR retrotransposons.

An important question now is: How do the piRNA and H3K9me2/3 findings relate to each other? Given the known effect of nuclear Piwi proteins in mouse and flies on chromatin, the most parsimonious explanation may be that the maternally-provided Ziwi-piRNA pool directs the H3K9 methylation, similar to what has been described for nuclear Piwi proteins in mouse and *Drosophila*. In the somatic cells and PGCs, this would prevent their expression – and in the PGCs this may additionally help define piRNA-generating loci. Indeed, such a maternal effect on piRNA clusters in the PGCs has been described in *Drosophila*^23^ and effects of maternal piRNAs in the soma of fly embryos have also been described^24^. However, we also find that the initial gain of H3K9me2/3 does not depend on transcription, a fact that does not match the current models on nuclear Piwi/Argonaute protein activities in eukaryotes which favors the targeting of the nascent transcripts, and not the DNA directly. This requires that any Ziwi-driven H3K9 methylation model to invoke a novel mode of target recognition, possibly involving Ziwi-piRNA complexes that recognize both DNA and RNA as target molecules. Here, we note that DNA targeting may potentially be facilitated by the serial DNA strand opening that occurs every 15 minutes via DNA replication during cleavage stage. In addition, antisense Ziwi-bound piRNAs and their biased inheritance/stability in PGCs may also enable them to conduct LTR sense transcript cleavage and/or silencing in PGCs to further limit the presence of functional TE-derived transcripts and safeguard the germline genome.

Our findings are also compatible with a model in which focal H3K9 methylation is driven by alternative factor(s) rather than Ziwi-piRNA complexes – however, these factor(s) would need to display the same specificity for LTRs as the maternal piRNA pool. A previous study demonstrated that H3K9me3 deposition at satellite repeats depends on transcription (miR-430 expression)^16^. However, our study suggests that the mechanism involved in initial LTR heterochromatinization is transcription independent. Here, KRAB-ZnF proteins would be excellent candidates for a transcription-independent heterochromatin trigger, since these factors act through direct DNA binding - if it were not for the fact KRAB-ZnF genes are absent from the zebrafish genome. Interestingly, the very same chromosome that harbors many of the young LTR elements targeted by piRNAs and H3K9 methylation, zebrafish chromosome 4, also harbors a large ZnF protein family of unknown function. However, transcripts for these ZnF family are not present in preZGA embryos ^25^ and the vast majority are transcribed at or after ZGA. These results, coupled with the fact that focal H3K9me2/3 acquisition at LTRs is transcription independent, would initially appear to limit their involvement. Here, future work will explore whether ZnF family proteins might be maternally inherited and help guide focal H3K9me2/3 silencing, either alone or physical in association with Ziwi-piRNA complexes – in addition to exploring a possible direct Ziwi-piRNA interaction with DNA.

## Methods

### Zebrafish husbandry and embryo collection

Zebrafish (*Danio rerio*) strains were maintained in accordance with approved institutional protocols at the University of Utah and at the IMP in Vienna, Austria. All experiments using zebrafish were approved by IACUC Protocol 20-04011 or by the ‘Amt der Wiener Landesregierung, Magistratsabteilung 58 - Wasserrecht’ (animal protocols GZ 342445/2016/12 and MA 58-221180-2021-16). Wildtype zebrafish were from the Tubingen (Tü) strain. For hybrid generation, TLAB fish, generated by crossing zebrafish AB and the natural variant TL (Tupfel Longfin) stocks, were used as wild-type zebrafish. Transgenic Tg(kop:EGFP-F-nanos1-3’UTR) and Tg(buc:GFP) strains were used for isolating primordial germ cells, and were kindly provided by Roland Dosch lab. Embryos were maintained at 28.5°C. All developmental staging were based on the time after fertilization and morphology confirmation as described^26^. Embryos for ChIP and FACS were dechorionated at 1-cell stage by pronase treatment and maintained in glass or agarose coated plastic petri dish until the desired stages.

### Medaka husbandry and embryo collection

Medaka (*Oryzias latipes*, CAB strain) were raised according to standard protocols (28°C water temperature; 14/10 hour light/dark cycle) and served as wild-type medaka. All medaka experiments were conducted according to Austrian and European guidelines for animal research and approved by the Amt der Wiener Landesregierung, Magistratsabteilung 58 – Wasserrecht under the protocol GZ: 198603/2018/14.

### Constructs and microinjection

Zebrafish codon-optimized KDM4E fragment was synthesized and cloned into pCS2+mCherry plasmid. RNAs were *in vitro* transcribed from linearized pCS2+-drKDM4E-mCherry and pCS2+-mCherry templates using mMESSAGE mMACHINE™ SP6 transcription kit (ThermoFisher Scientific, AM1340), followed by poly adenylation reaction using Poly(A) tailing kit (ThermoFisher Scientific, AM1350). 200pg KDM4E-mCherry mRNA or 100pg mCherry mRNA was injected into 1-cell zebrafish embryos. Live embryos were collected at 4 hpf and 5.3 hpf. For each batch, 25-30 embryos were used for western blotting and the 30-50 embryos were snap frozen for RNA extraction. Three to four biological replicates were collected for each condition.

### Drug treatment

1-cell zebrafish embryos were injected with 0.2ng α-amanitin as described^8^. Live embryos were collected at 4 hpf and 4.5 hpf according to the development of uninjected control since treated embryos arrest around 4 hpf. A-amanitin treated embryos were used for western blotting and whole mount staining. 1-cell zebrafish embryos were dechorionated and treated with 1.5μM flavopiridol as described^27^. Live embryos were collected at 4 hpf for ChIP.

### Western blotting

4 hpf and 4.5 hpf zebrafish embryos were dechorionated by pronase and washed with 1X PBS twice. Embryo yolks were removed by adding 3X deyolking buffer^28^. Cells were pelleted and lysed with strong cell lysis buffer (10mM Tris-HCl pH=7.5, 50nM NaCl, 1% Triton, 0.1% SDS and 1X proteinase inhibitor) on ice for 10min. Samples are boiled at 95°C for 5min after 4X sample buffer added. Western blotting was performed as standard procedure. H3K9me2 (Abcam 1220, 1:1000), H3K9me3 (Active Motif 39161, 1:1000) and H3 antibodies (Abcam ab1791 and Active motif 39763, 1:10000) were used. Signal was acquired and quantified using LI-COR Odyssey imaging system.

### Embryo whole-mount staining and imaging

Zebrafish whole mount staining was performed as described^29^. H3K9me2 (Abcam 1220) and H3K9me3 (Active Motif 39161) antibody were used as primary antibodies. Donkey anti-Rabbit IgG Alexa Fluor 488 (Invitrogen, A-21206) and donkey anti-mouse IgG Alexa Fluor 594 (Invitrogen, A-21203) were used as secondary antibodies.

### Ziwi-IP

50-100 embryos/oocytes were lysed in 600 μl lysis buffer (25mM Tris-Cl pH7.5, 150mM NaCl, 1.5mM MgCl2, 1% Triton, protease inhibitors, 1mM DTT, antiRNase 1:1000), crushed by a pestle and sonicated on bioruptor (5 cycles 30’’on/30’’off on high power). Anti-Ziwi antibody (homemade, 1:200 dilution) was added to the cell lysate followed by 2h rotation at 4°C, then 30μl Protein G dynabeads was added and rotated at 4°C for 1h. Beads were washed 2 times with wash buffer (50mM Tris-Cl pH 7.5, 1M NaCl, 1mM EDTA, 1% Igepal CA-630, 0.1% SDS, 0.5% sodium deoxycholate) and put in 500μl Trizol. RNA extraction was performed following Trizol protocol, additional sodium acetate was added and precipitated overnight with glycoblue at −20°C. RNAs were labeled with P^32^ (all IP RNA in 10μl with 5μCi gamma-ATP), and run on a 15% denaturing gel.

### FACS

Tg(buc:GFP) inbreed or outbreed (Tg(buc:GFP) female x WT Tü male) embryos were collected at 2.5 hpf and 4 hpf. Tg(kop:EGFP-F-nanos1-3’UTR) inbreed embryos were collected at 12 hpf and 24 hpf. For 2.5 hpf embryos, ~500 dechorionated embryos were washed twice and resuspended in pre-cold 5% FBS-PBS with 0.5M sucrose to prevent cell bursting. Cell resuspension was filtered through 100μm cell strainer and kept on ice. For 4hpf embryos, dechorionated embryos were washed three time with Ca^2+^-free Ringer solution to remove yolk and resuspended in 5% FBS-PBS. Cell resuspension was filtered through 40μm cell strainer. For 12 hpf and 24 hpf embryos, dechorionated embryos were washed three times with Ca^2+^-free Ringer solution, and digested with pre-warmed trypsin at 37°C for 30min. FBS was added at 1:10 ratio and 1M CaCl2 was added at 1:1000 ratio. Cells were resuspended in 5% FBS-PBS and filtered through 40μm cell strainer. DAPI (1:1000) was added to all samples before sorting.

Cell sorting was performed on BD FACSAria with 130μm nozzle for 2.5 hpf embryos, 100μm nozzle for 4 hpf embryos and 85μm nozzle for 12 hpf and 24 hpf embryos. Cells for sRNA-seq were directly collected in RLT buffer and cells for ATAC-seq were collected in 5% FBS-PBS. Two or three biological replicates of each cell types at each developmental stage were collected.

### Hybrid embryo generation

Zebrafish hybrid and medaka hybrid embryos were generated as described before ^21,22^. Three or four biological replicates were collected at each developmental stage for each type of hybrid embryos.

### RNA extraction

30-50 zebrafish embryos were snap frozen at desired developmental stage for each biological replicate. RNA extractions were performed using AllPrep DNA/RNA mini kit (Qiagen, 80004) with modifications. RWT buffer was used to recover both small (<200nt) and long RNAs. Extracted total RNAs were treated with TURBO DNA-free kit (Invitrogen, AM1907) to remove residual DNAs. For FACS samples, 1000-5000 PGCs or 10000 somatic cells were sorted into RLT buffer. Total RNAs were extracted using AllPrep DNA/RNA Micro Kit (Qiagen, 80204). For hybrid and medaka purebred embryos, 10-50 embryos were collected in Trizol. Total RNAs were extracted using Qiagen miRNeasy kit (Qiagen, 217004).

### RNA Library preparation and sequencing

Sample quality were evaluated by Tapestation and only samples with RIN>8.0 (7.0 for hybrid embryos) were proceed to library preparation. Three biological replicates were collected for each developmental stage unless indicated.

Ribosomal RNA depletion from total RNA was performed using either the RiboCop rRNA Depletion Kit (Lexogen) using 500 ng of RNA per sample (zebrafish, zebrafish hybrid and medaka samples) or the Ribo-zero gold kit (medaka hybrid and mCherry or KDM4E-injected zebrafish samples). Libraries were prepared using NEBNext Ultra Directional RNA Library Prep Kit for Illumina (NEB) (zebrafish, zebrafish hybrid and medaka samples) or the Illumina TruSeq Stranded Total RNA Library Prep kit (medaka hybrid and mCherry or KDM4E-injected zebrafish samples) and checked using a Fragment Analyzer System (Agilent) before multiplexing. Sequencing was performed on Illumina HiSeqV4 SR100 and NovaSeq 150 bp paired-end sequencing platform.’

Small RNA libraries for WT zebrafish, medaka, hybrid embryos and sorted cells were prepared using QIAseq miRNA Library Prep with single index or UDI kit (Qiagen, 331502) and sequenced on HiSeq 65bp single-end sequencing or NovaSeq 50bp paired-end sequencing platform. sRNA libraries for *dicer-null* embryos were prepared using Illumina TruSeq Small RNA Sample Prep kit (Illumina, RS-200-0012) and sequenced on HiSeq 50bp single-end sequencing platform. sRNA libraries for Ziwi-IP samples were prepared using NEBNext Multiplex Small RNA Library Prep Set for Illumina (NEB, E7300S) and sequenced on HiSeq 50bp single-end sequencing platform.

### Chromatin immunoprecipitation and library prep

200 dechorionated embryos were collected in 1X PBS and fixed with 1% formaldehyde at room temperature for 10min. 125mM glycine was added to quench the fixation for 5min. Embryos were washed with cold PBS twice. Samples were snap frozen and stored at −80°C until enough samples were collected for ChIP experiments. 4500 2.5 hpf embryos or 1000 4 hpf embryos were used for each replicate, two or three biological replicates were collected for each condition.

1000 embryos were lysed in 1ml mild cell lysis buffer (10mM Tris-HCl pH=8.0, 10mM NaCl, 0.5% NP-40, 1x proteinase inhibitor) for 10min at 4°C. Nuclei were pelleted and washed with 1ml nuclei wash buffer (50mM Tris-HCl pH=8.0, 100mM NaCl, 10mM EDTA) at room temperature for 10min twice followed by incubation with 100μl nuclei lysis buffer (50mM Tris-HCl pH=8.0, 10mM EDTA, 1% SDS) on ice for 10min. 900μl IP dilution buffer (0.01% SDS, 1.1% Triton X-100, 1.2mM EDTA, 16.7ml Tris-HCl pH=8.0, 167mM NaCl) was added. DNA was sonicated to 200-700bp with Branson sonifier. About 1ml soluble chromatin was pre-cleared with 20μl Ampure beads for 1h at 4°C, and then 100μl samples was taken as input. 5μg of H3K9me2 (Abcam 1220) or H3K9me3 (Abcam 8898) antibody was added to the chromatin followed by overnight incubation at 4°C. To pull down the antibody associated chromatin, 50μl Ampure beads were added and rotated at 4°C for 4-6h. Beads were washed with RIPA buffer (10mM Tris-HCl Ph=7.5, 140mM NaCl, 1mM EDTA, 0.5mM EGTA, 1% Triton X-100, 0.1% SDS, 0.1% Sodium deoxycholate) eight times, LiCl washing buffer (10mM Tris-HCl pH=8.0, 1mM EDTA, 150mM LiCl, 0.5% NP-40, 0.5% DOC) twice and TE buffer (10mM Tris-HCl pH=8.0, 1mM EDTA) twice. 100ul elution buffer (10mM Tris-HCl pH=8.0, 5mM EDTA, 300mM NaCl, 0.1% SDS) was added to the beads for IP samples and reverse crosslinked together with input sample at 65°C for 6h. 20mg of RNase was added and incubated at 37°C for 30min. 5μl Proteinase K was added and incubated at 50°C for 2h. DNA was extracted with Ampure beads and eluted with 40μl 10mM Tris-HCl pH=8.0.

ChIP-seq libraries were prepared with NEBNext ChIP-Seq Library Prep Master Mix (NEB, E6240L) and then amplified with NEBNext High-Fidelity 2X PCR Master Mix (NEB, M0541L). Libraries were sequenced on HiSeq 125bp paired-end platform.

### ATAC and library preparation

Sorted PGCs or somatic cells were lysed with 4μl ice-cold 1x lysis buffer (10mM Tris-HCl pH=7.4, 3mM MgCl_2_, 0.1% NP-40) on ice for 15min. 5μl H_2_O, 12.5ul 2X Tris-DMF-tagmentation buffer (20mM Tris-HCl pH=8.0, 10mM MgCl_2_, 20% DMF) and 2.5ul Nextera Tn5 transposase were added to the lysed cells. The reaction was incubated at 37°C for 30min. 0.5μl 10% SDS was added to stop the reaction followed by 5min incubation on ice. Tagmented DNAs were purified with Qiagen MinElute PCR purification kit (Qiagen, 28006) and eluted with 22μl EB buffer.

ATAC-seq library was amplified from 20μl tagmented DNA with 25μl 2x Phusion PCR master mix (NEB, 28006) and 0.625μl of each 100mM library primers. Five cycles of pre-amplification PCR were performed (72°C for 5min, 5 cycles of 95°C 30s and 72°C 90s, hold at 4°C). 5μl of pre-amp product was used for qPCR with the same cycling condition to determine the number of additional PCR cycles required for library prep. Proper additional PCR cycles were done for the rest of pre-amp reaction and finished with 10min incubation at 72°C. Library was purified using Ampure beads and checked by Tapestation. ATAC-seq libraries were sequenced on HiSeq 125bp paired-end platform or Novaseq 50bp paired-end platform.

### *De novo* transposon annotation

*De novo* transposable element annotation for medaka (ensemble 90 release) genome was completed using EDTA pipeline ^30^with following parameters: EDTA.pl --overwrite 0 --genome OL_ens90_sim.fa --species others --step all --cds OL_cds_ens90.fa --sensitive 1 --anno 1 --evaluate 1. LTR annotations for medaka genome were combined with zebrafish LTR annotations from repeatmasker for the following analyses.

### sRNA-seq data analysis

Unique molecular index (UMI) was firstly extracted from the paried-end fastq files of Qiagen sRNA libraries using script smallRNA_pe_umi_extractor.pl. The generated fastq files were aligned to Zv10 zebrafish genome assembly using STAR (https://github.com/alexdobin/STAR) with the following parameters: STAR --outSAMtype BAM SortedByCoordinate --outFilterMismatchNoverLmax 0.05 --outFilterMatchNmin 16 --outFilterScoreMinOverLread 0 --outFilterMatchNminOverLread 0 --outFilterMultimapNmax 400 --winAnchorMultimapNmax 800 --seedSearchStartLmax 13 --alignIntronMax 1 --runRNGseed 42 --outMultimapperOrder Random --outSAMmultNmax 1. Reads that map to multiple locations with equal mapping quality score will be assign to one location randomly in the genome. Duplicated reads were removed from Bam files using script bam_umi_dedup.pl. Single-end fastq files of illumina or NEBNext libraries were directly aligned to the genome using STAR with the same parameters. sRNA fastq files from hybrid embryos and corresponding purebred controls were processed in the same way except aligning to zebrafish and medaka combined genome.

Read size and sequence frequency information were extracted using get_bam_seq_stats.pl script from biotoolbox pacakge (https://github.com/tjparnell/biotoolbox). Sequence logo were generated using ggplot2 (https://ggplot2.tidyverse.org/) and ggseqlogo (https://github.com/omarwagih/ggseqlogo) packages in R. Size selection was accomplished by filter_bam.pl script from biotoolbox. Genomic distribution of sRNAs was determined by counting reads intersect with zebrafish repeatmasker annotations from UCSC using featureCounts (http://subread.sourceforge.net/). LTR targeted antisense/sense piRNAs were counted in regard to the orientation of LTR annotation using featureCounts. Ping-pong signature were generated for LTR targeted piRNAs using small_rna_signature package (https://github.com/ARTbio/tools-artbio/tree/master/tools/small_rna_signatures).

### Total RNA-seq analysis

Adapters from paired-end RNA-seq fastq files were removed using Cutadapt (https://cutadapt.readthedocs.io/en/stable/) and then aligned to the Zv10 zebrafish genome assembly using STAR with the following parameters: --twopassMode Basic --outSAMtype BAM SortedByCoordinate --quantMode TranscriptomeSAM --outWigType bedGraph --outWigStrand Unstranded --outFilterMultimapNmax 100 --winAnchorMultimapNmax 200 --outMultimapperOrder Random --runRNGseed 42 --outSAMmultNmax 1. Multimapped reads were assigned to one location in the genome randomly. All RNA-seq fastq files from hybrid embryos and corresponding purebred controls were aligned to a combined zebrafish and medaka genome index (both from ensembl 90 release). DR (*Danio rerio*) and OL (*Oryzias latipes*) were added to the chromosome name to distinguish the origins.

Data for metaplot was collected using get_binned_data.pl from biotoolbox and plotted by ggplot2 in R. LTR counts and gene counts were obtained by featureCounts and combined for differential expression analysis by DESeq2 (https://bioconductor.org/packages/release/bioc/html/DESeq2.html) in R (p<0.05, log2FC>=1).

### ChIP-seq data analysis

Paired-end fastq files were aligned to Zv10 zebrafish genome assembly using Novoalign (http://www.novocraft.com/products/novoalign/) with the following parameters: -o SAM -r Random. Multimappers were assigned to one location in the genome randomly. ChIP-seq peaks were called by integrating multiple replicates using Multi-Replica Macs ChIPSeq Wrapper (https://github.com/HuntsmanCancerInstitute/MultiRepMacsChIPSeq) with the following parameters: --pe --nodup --cutoff 2. Heatmap was generated using computeMatrix reference-point and plotHeatmap in deepTools package (https://deeptools.readthedocs.io/en/develop/content/list_of_tools.html). ChIP-seq signal tracks (log2 fold enrichment) were generated by Multi-Replica Macs ChIPSeq Wrapper and visualized in IGV genome browser (https://software.broadinstitute.org/software/igv/).

### ATAC-seq data analysis

Paired-end fastq files were aligned to Zv10 zebrafish genome assembly using Novoalign with the following parameters: -o SAM -r Random. Reads mapped to mitochondria were removed using the following command: sed ‘/chrM/d’ <atac_random.sam> atac_random_noChrM.sam and converted to Bam files using Samtools (http://www.htslib.org/). Chromatin accessibility were determined by integrating multiple replicates using Multi-Replica Macs ChIPSeq Wrapper with the following parameters: --pe --min 30 --max 120 --nodup --cutoff 2.

### Data access

High-throughput sequencing data from this study were deposited at Gene Expression Omnibus under accession number GSExxxx.

### Reanalyzed public datasets

Datasets are available under the following accession numbers: PGCs and somatic cells RNA-seq: GSE122480

## Author Contributions

Y.G. performed all experiments and data analyses unless indicated. K.R.G. generated hybrid embryos and collected purebred embryos. S.L. performed Ziwi-IP and collected *dicer null* embryos. M.E.P piloted this project. C.L.W assisted in optimizing KDM4E injection. E.J.G assited in experimental design. B.R.C., A.P., and R.F.K. supervised the project. Y.G. and B.R.C. prepared the manuscript with input from all authors.

## Acknowledgement

We thank J. Guo, C. Yi, N. Verma, S. Shadle and all other members in the Cairns laboratory for fruitful discussions, helpful bioinformatic and technical expertise; W. Xie lab and F. Muller lab for sharing the experimental protocols; R. Dosh lab for gifting us the transgenic zebrafish strains; M. Hobbs, S. Nielsen, R. Stewart, G. Nikkum, C. James, J. Lee, L. Graham for assistance with animal husbandry. Special thanks to B. Dalley for sequencing expertise, T. Parnell for bioinformatic expertise, J. Marvin, T. Galland and N. Choksi for flow cytometry expertise. Funding for this work involved Howard Hughes Medical Institute (support of B. R. Cairns); FFG (Headquarter grant FFG-852936), the FWF START program (Y 1031-B28), the HFSP Career Development Award (CDA00066/2015), the HFSP Young Investigator Award, and EMBO-YIP funds to A. Pauli; a DOC fellowship from the Austrian Academy of Sciences (OeAW) to K.R. Gert. Sample sequencing was performed at the High-Throughput Genomics and Bioinformatic Analysis Shared Resources (NCI, P30CA042014) at Huntsman Cancer Institute. Imaging was performed at the University of Utah Cell Imaging Core (1S10RR024761-01). Flow cytometry was performed at University of Utah Flow Cytometry Facility (NCI, 5P30CA042014-24; NIH, 1S10RR026802-01). Animals were maintained in University of Utah Centralized Zebrafish Animal Resource (CZAR) facility (NIH, 1G20OD018369-01).

