## Supplemental Figures and Legends for "Maternally-inherited anti-sense piRNAs antagonize transposon expression in teleost embryos"

Supp. Figure 1

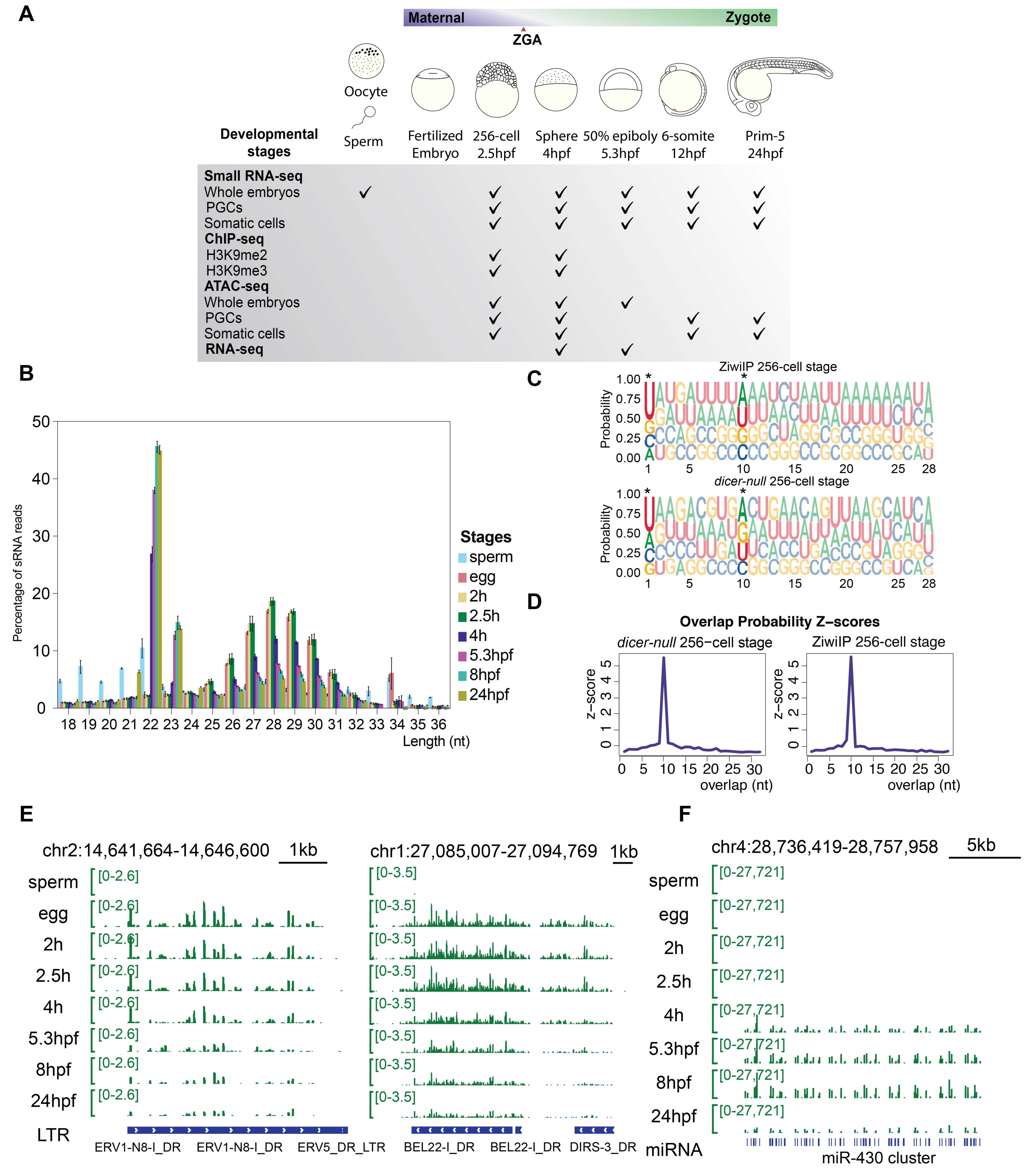


**Figure S1. Analyses of small RNA populations in zebrafish gametes and early embryos, related to Figure 1. (A)** Datasets are generated for this study, including sRNA-seq, total RNA-seq, ChIP-seq and ATAC-seq. **(B)** sRNA size distribution in zebrafish egg, sperm and early embryos. piRNA-sized sRNAs (24-32nt) are the most abundant type of sRNAs in egg and preZGA embryos (2 hpf and 2.5 hpf), and gradually decrease after ZGA (4 hpf, 5.3 hpf, 8hpf and 24 hpf). MiRNAs (22-23nt) abundance increases after ZGA. **(C)** sRNAs from Ziwi-IP and *dicer-null* embryos carry piRNA sequence bias: U1 and A10 (highlighted and marked by *). **(D)** sRNAs from Ziwi-IP and *dicer-null* embryos show piRNA ping-pong signature (10bp overlap). **(E)** Genome browser snapshots of representative LTR-targeting piRNAs in zebrafish egg and early embryos. **(F)** Genome browser snapshot of miR-430 cluster, which is known to activate during ZGA.

### Supp. Figure 2

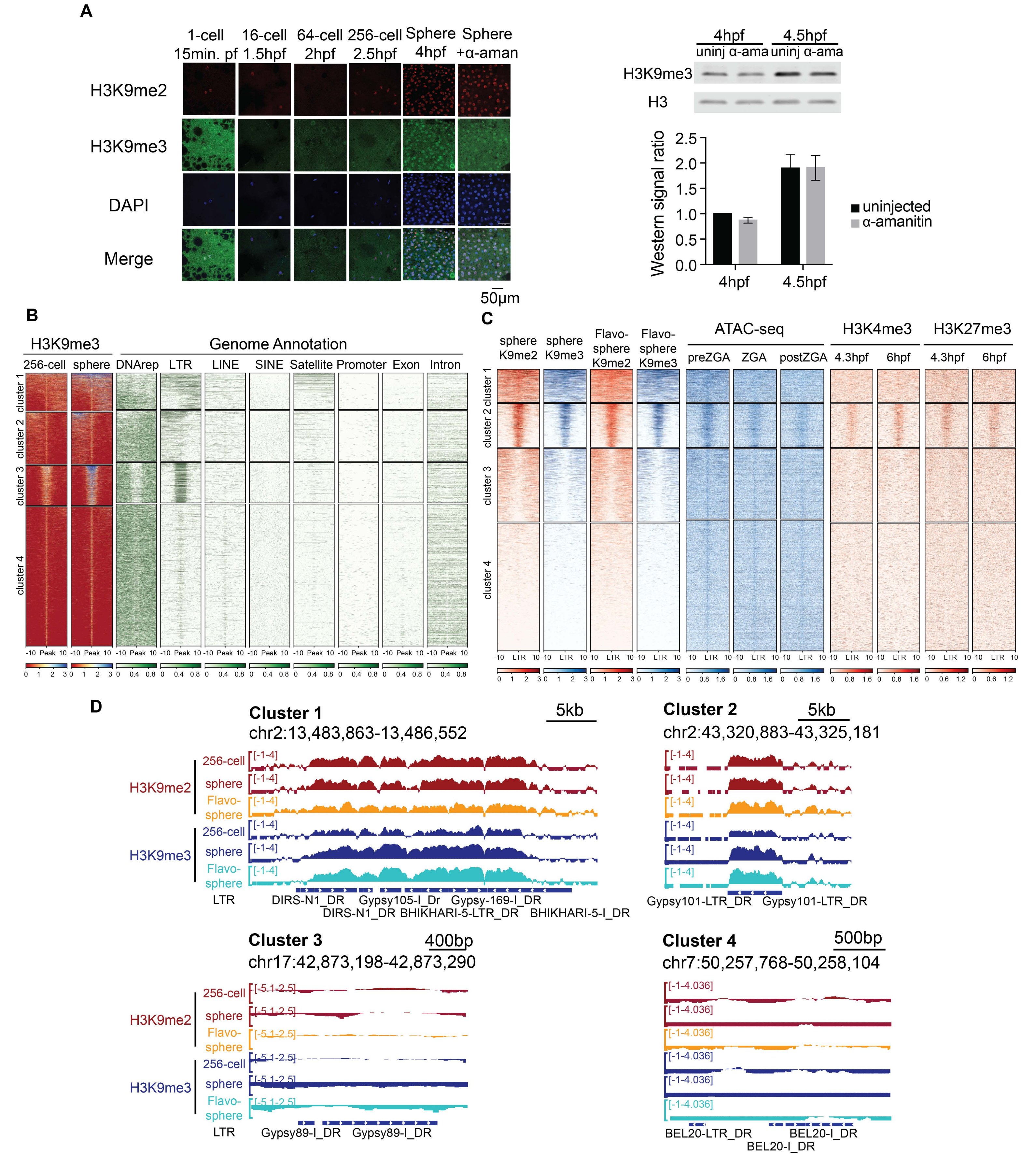


**Figure S2. The repressive histone mark H3K9me3 is established at evolutionarily young LTRs in a transcription-independent manner, related to Figure 2. (A)** Left: Immunofluorescent staining of H3K9me2 (red) and H3K9me3 (green) shows the dynamics of heterochromatin during early embryogenesis. Right: western blotting shows transcription inhibition treatment (α-amanitin) doesn't affect the establishment of H3K9me2/3. **(B)** H3K9me3 ChIP peaks are enriched at Satellite repeats (cluster 1) and LTRs (clusters 1, 2 and 3) in zebrafish genome. **(C)** Heatmap of H3K9me2 and H3K9me3 ChIP with and without the transcription inhibitor flavopiridol treatment (1.5uM) at sphere stage, ATAC-seq, H3K4me3 and H3K27me3 ChIP at LTR internal region clusters. **(D)** Genome browser snapshots of H3K9me2 and H3K9me3 at representative LTR internal regions. Clusters in panel C and D are defined in Figure 2C.

Supp. Figure 3

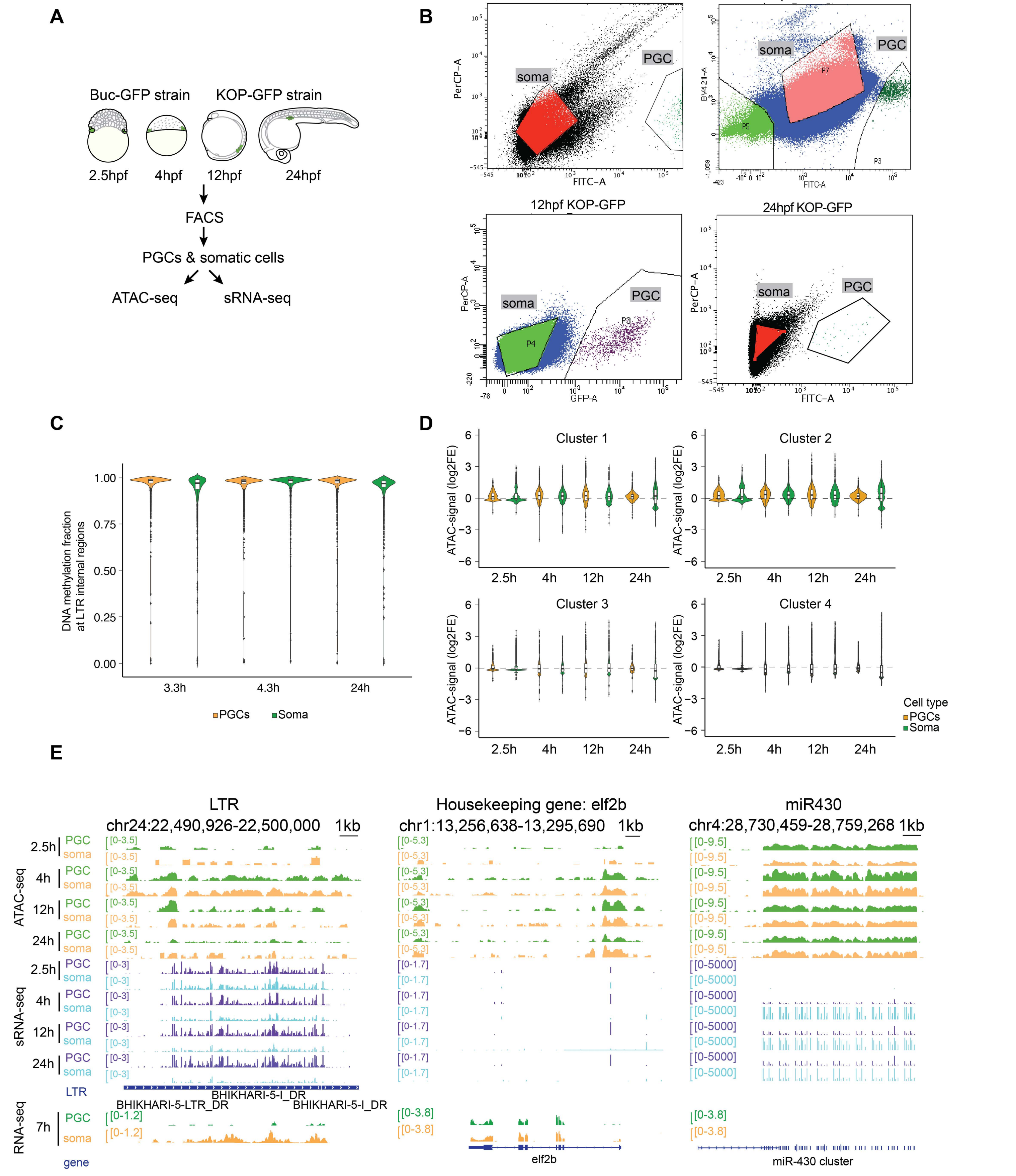


**Figure S3. The chromatin landscape shows limited differences in PGC and somatic cells, related to Figure 3. (A)** Schematic representation of the experimental setup of obtaining and sequencing PGCs and somatic cells from zebrafish early embryos. **(B)** PGCs and somatic cell isolation from early embryos via flow cytometry. **(C)** Reduced representative bisulfite sequencing data shows DNA are hypermethylated at LTRs in PGCs and somatic cells. **(D)** Chromatin accessibility of LTR internal region clusters in PGCs and soma has no significant difference. Clusters are the same as Figure 2C. **(E)** Genome browser snapshots of chromatin accessibility, sRNAs and total RNA abundance at LTR, housekeeping gene and miR-430 locus.

**Supp. Figure 4**

**A**

**Repeats Annotation for Medaka Genome by EDTA pipeline**
Repeat Classes
Total Sequences: 6751
Total Length: 700386597 bp

| Class | Count | bpMasked | %masked |
|---|---|---|---|
| **DNA** | - | - | - |
| DTA | 224574 | 55398922 | 7.91% |
| DTC | 251591 | 43894316 | 6.27% |
| DTH | 45192 | 9414487 | 1.34% |
| DTM | 188203 | 33919119 | 4.84% |
| DTT | 16781 | 2412145 | 0.34% |
| Helitron | 77283 | 14062487 | 2.01% |
| **LTR** | - | - | - |
| Copia | 2518 | 304284 | 0.04% |
| Gypsy | 27940 | 7942045 | 1.13% |
| unknown | 59612 | 9888872 | 1.41% |
| **MITE** | - | - | - |
| DTA | 26842 | 4720495 | 0.67% |
| DTC | 7107 | 1048400 | 0.15% |
| DTH | 1778 | 317163 | 0.05% |
| DTM | 5339 | 708051 | 0.10% |
| DTT | 263 | 26905 | 0.00% |
| **Total** | 935023 | 184057691 | 26.28% |

**B**

RNAseq data from zebrafish, medaka, zebrafish hybrid and medaka hybrid

↓

Align to zebrafish+medaka combined genome by STAR (multimappers were assigned to one location randomly)

↓

Obtain read counts for genes and LTRs

↓

Identify activated genes and LTRs using DESeq2 (log2FC>1, p<0.05)

**C**

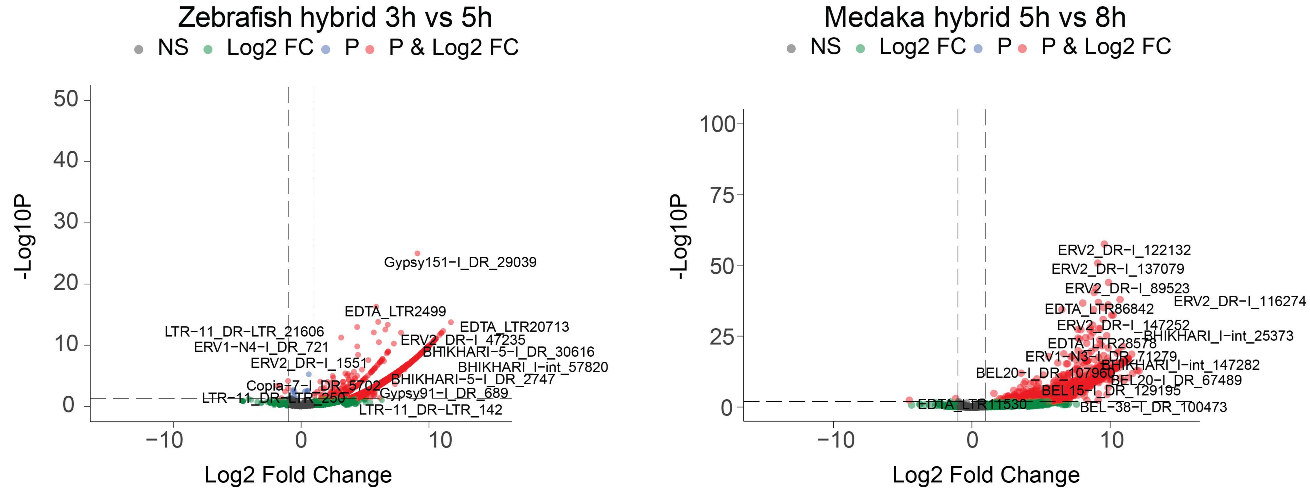


**D**

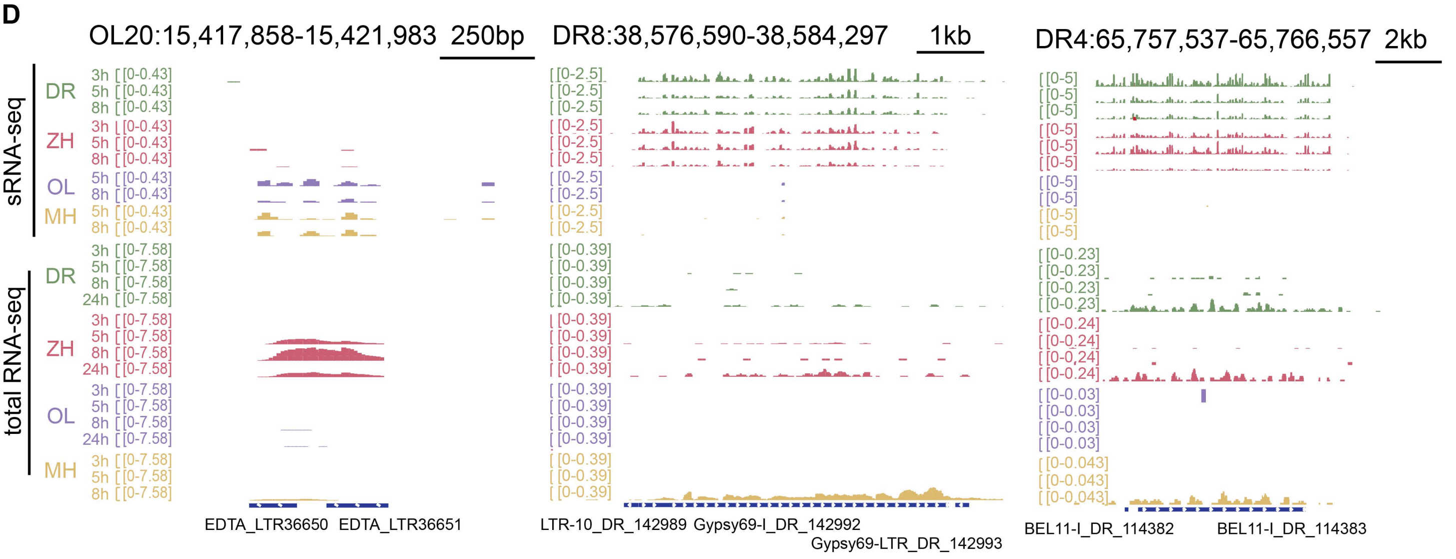


**Figure S4. LTRs are desilenced in the absence of targeting piRNAs, related to Figures 4 and 5. (A)** A summary of de novo annotated transposons in medaka genome by the EDTA pipeline. **(B)** The workflow of identifying activated genes and LTRs from zebrafish, medaka, zebrafish hybrid and medaka hybrid embryos. **(C)** Volcano plots of differentially expressed LTRs in zebrafish hybrid (3h vs 5h, left) and medaka hybrid (5h vs 8h, right) embryos ($p<0.05$, log2FC>=1). **(D)** Genome browser snapshots of representative activated medaka and zebrafish LTRs in hybrid embryos (zebrafish, DR; medaka, OL, zebrafish hybrid, ZH; medaka hybrid, MH).
